# Sex reversal and ontogeny under climate change and chemical pollution: are there interactions between the effects of elevated temperature and a xenoestrogen on early development in agile frogs?

**DOI:** 10.1101/2020.12.29.424761

**Authors:** Zsanett Mikó, Edina Nemesházi, Nikolett Ujhegyi, Viktória Verebélyi, János Ujszegi, Andrea Kásler, Réka Bertalan, Nóra Vili, Zoltán Gál, Orsolya I. Hoffmann, Attila Hettyey, Veronika Bókony

## Abstract

Anthropogenic environmental change poses a special threat to species in which genetic sex determination can be overwritten by the thermal and chemical environment. Endocrine disrupting chemicals as well as extreme temperatures can induce sex reversal in such species, with wide-ranging consequences for fitness, demography, population viability and evolution. Despite accumulating evidence suggesting that chemical and thermal effects may interact in ecological contexts, little is known about their combined effects on sex reversal. Here we assessed the simultaneous effects of high temperature (masculinizing agent) and 17α-ethinylestradiol (EE2), a widespread xenoestrogen (feminizing agent), on sexual development and fitness-related traits in agile frogs (*Rana dalmatina*). We exposed tadpoles to a six-days heat wave (30 °C) and/or an ecologically relevant concentration of EE2 (30 ng/L) in one of three consecutive larval periods, and diagnosed sex reversals two months after metamorphosis using species-specific markers for genetic sexing. We found that high temperature induced female-to-male sex reversal, decreased survival, delayed metamorphosis, decreased body mass at metamorphosis, and increased the proportion of animals that had no fat bodies, while EE2 had no effect on these traits. Simultaneous exposure to heat and EE2 had non-additive effects on juvenile body mass, which were dependent on treatment timing and further complicated by a negative effect of sex reversal on body mass. These results show that environmentally relevant exposure to EE2 does not diminish the masculinizing effects of high temperature. Instead, our findings on growth suggest that climate change and chemical pollution may have complex consequences for individual fitness and population persistence in species with environment-sensitive sex determination.

## 1. Introduction

Our ecosystem is facing a multitude of rapid environmental changes induced by human activities, including climate change and various forms of environmental pollution. These anthropogenic environmental changes can affect the neuroendocrine, cardiorespiratory, iono- and osmoregulatory, immune and reproductive systems of wild animals influencing their growth and development especially during early life (Bernanke and Köhler, 2009; Burraco et al., 2020; Parmesan, 2006; Sparling et al., 2010; Whitney et al., 2016). Moreover, both climate change and environmental contaminants pose a special threat to species in which sex determination and sexual development are sensitive to environmental conditions. For example, sex determination is purely temperature-dependent in several ectothermic vertebrates, thus climate change puts such species in danger of skewed sex ratios, with potentially wide-ranging consequences from evolutionary changes in breeding systems and social behaviors (Liker et al., 2014, 2013) and reduced genetic diversity to demographic collapse (Mitchell and Janzen, 2010). Furthermore, in several species of fish, amphibians, and reptiles, sex is determined genetically but is additionally influenced by thermal or chemical stimuli experienced during early ontogeny, which can result in sex reversal, a mismatch between genetic sex and phenotypic sex (Eggert, 2004; Holleley et al., 2016; Ospina-Alvarez and Piferrer, 2008). The consequences of sex reversal go beyond skewed sex ratios in these species as well, because sex-reversed individuals may have lower or higher reproductive success than conspecifics in which phenotypic sex and chromosomal sex are concordant (Holleley et al., 2016; Senior et al., 2012), and may produce sex-biased offspring which may lead to changes in demography, sex-chromosome evolution and even to extinction (Bókony et al., 2017; Nemesházi et al., 2020b; Quinn et al., 2011; Schwanz et al., 2020; Wedekind, 2017). Despite potentially widespread occurrence of sex reversal and its diverse implications, this phenomenon is rarely studied, mainly because the identification of genetic sex requires species-specific molecular methods in ectothermic vertebrates (Nemesházi et al., 2020a). Nevertheless, sex ratios can also be biased by sex-specific tolerances to environmental stress (Afonso et al., 2003; Liwanag et al., 2018; Medina et al., 2002), hence it is important to distinguish between environmental effects on sex-specific mortality *versus* on sex determination (Geffroy and Wedekind, 2020).

Exposure to high temperatures during the sensitive period of sexual development can result in both masculinization and feminization in reptiles, depending on the species (Holleley et al., 2016; Mitchell and Janzen, 2010), whereas in amphibians and fish, heat usually leads to the development of genetically female individuals into phenotypic males via the disruption of the enzymatic machinery of estrogen synthesis (Baroiller and D’Cotta, 2016; Eggert, 2004; Lambert et al., 2018). In taxa prone to heat-induced masculinization, female-to-male sex reversal is expected to become more frequent as a result of global climate change, which includes both rises in average temperatures and an increasing frequency of extreme weather events such as heat waves (IPCC, 2014; Spinoni et al., 2015; Tomczyk and Bednorz, 2019). Water temperature in ponds, where the young of many aquatic vertebrates develop, can already reach as high as 30-50 °C during spring and summer, even under temperate and highland Mediterranean climates (Lambert et al., 2018; Lindauer et al., 2020), and heat waves may have particularly strong effects on populations living in urbanized areas because of the urban heat island effect (Brans et al., 2018). Accordingly, trends observed in phenotypic sex ratios suggest that the expected increase in heat-induced masculinization rates has already started in some species (Bókony et al., 2017; Grayson et al., 2014).

In addition to climatic challenges, several pollutants that are released into the environment act as endocrine disruptor chemicals (EDCs), which can affect somatic and sexual development by interfering with the hormonal system of animals (Orton and Tyler, 2015). Various EDCs, such as artificial hormones (e.g. contraceptives, growth promoters) and other substances (e.g. pesticides) can cause abnormalities in gonadal development including complete sex reversal (Flament, 2016; Orton and Tyler, 2015) and intersex, a form of incomplete sex reversal where male and female tissues occur simultaneously in the gonads (Abdel-Moneim et al., 2015; Ujhegyi and Bókony, 2020). The interactions (i.e. non-additive effects) between these two major environmental factors, i.e. temperature and EDCs, are being investigated with increasing intensity (Noyes and Lema, 2015), as climate change may modify the sensitivity of organisms to EDCs, while the chemicals may damage the capacity of organisms to respond to rapidly changing climatic conditions (Hooper et al., 2013; Noyes and Lema, 2015). However, very little is known about the combined effects of heat and EDCs on sex reversal, specifically. Results from experiments on fish showed that high temperature amplified the effect of clotrimazole, a masculinizing EDC present in many fungicides (Brown et al., 2015), while treatment with the female sex hormone 17β-estradiol, a natural hormone used in medication for menopausal symptoms, completely neutralized the masculinizing effect of high temperatures (Kitano et al., 2012, 2007). However, these studies used high concentrations of EDCs, which exceeded environmentally relevant concentrations by an order of magnitude.

In natural waters, hormonally active chemical agents typically occur in low concentrations (Loos et al., 2009), although many of the more persistent EDCs may be enriched in shallow, small water bodies and in their sediments (Bókony et al., 2018). 17α-ethinylestradiol (EE2) is a synthetic female sex hormone used in many contraceptive formulations, hence significant amounts are excreted with urine into wastewater. With the conventional wastewater treatments it cannot be removed completely, so it is often found in surface waters and ground waters worldwide, typically in concentrations of a few ng/L (Bhandari et al., 2015). In vertebrates, EE2 has a strong feminizing effect (Bhandari et al., 2015; Tamschick et al., 2016), but sensitivity differs among species, with effective concentrations varying from as low as 1.8 ng/L (Berg et al., 2009; Gyllenhammar et al., 2009) to as high as 500 ng/L (Tamschick et al., 2016) or even 1 µg/L (Mackenzie et al., 2003). It is unclear if the low concentrations of EE2 that occur in nature can compensate for the masculinizing effect of high temperatures that occur during heat waves, and if there are any other interactions between the effects of heat and EE2 that may influence the ecological consequences of climate change in polluted waters.

In this study, our aim was to investigate the simultaneous effects of high temperature and EE2 on the sexual development of the agile frog (*Rana dalmatina*), utilizing a molecular marker set recently developed for diagnosing sex reversals in this species (Nemesházi et al., 2020a). The agile frog is widespread in Europe, inhabiting natural woodlands as well as urbanized areas, and its population trends are decreasing (Kaya et al., 2009). We performed an experiment to mimic environmentally realistic scenarios in which we exposed tadpoles for a few days to high temperature (simulating a heat wave), EE2 (simulating a short-duration point source pollution), or both. We chose a temperature and EE2 concentration that are relatively high within the range of contemporary environmental conditions, but can be expected to occur with increasing frequency in nature because of climate change (Spinoni et al., 2015; Tomczyk and Bednorz, 2019) and ongoing urbanization. Furthermore, because both high temperature and EE2 are known to influence mortality, growth, and somatic development during early ontogeny in aquatic vertebrates (Gyllenhammar et al., 2009; Hogan et al., 2008; Hu et al., 2019; Marques da Cunha et al., 2019; Tompsett et al., 2012), we also investigated whether there is interaction between the effects of high temperature and EE2 on these fitness-related traits besides sexual development.

## 2. Materials and methods

### 2.1. Experimental procedures

On 8 March 2019 we collected 60 agile frog eggs from each of four freshly laid egg masses from three ponds (Apátkút, 47°46□28□N, 18°59□10.5□E; Ilona-tó, 47°42□47.7□N, 19°02□25.8□E; and Katlan, 47°42□40□N, 19°02□44.5□E) located in a hilly woodland in Hungary. Eggs were transported to the Experimental Station of the Plant Protection Institute (Centre for Agricultural Research) in Julianna-major, Budapest (47°32□52□N, 18°56□05□E). Until hatching (17 March), we kept eggs at 16.3 ± 0.3 °C (mean ± SD), each of the twelve clutches (families hereafter) separately in 5-L containers (24 × 16 ×13 cm) filled with 1.3 L reconstituted soft water (RSW, APHA 1985; Bókony et al., 2020). To ensure sufficient oxygenation, we aerated the water in the containers with aquarium air pumps. When the hatchlings reached the free swimming state (developmental stage 25 according to Gosner, 1960) four days after hatching, we started the experiment by randomly selecting 48 healthy-looking larvae from each of the 12 families and placing them into individual rearing containers. The remaining eggs and tadpoles were released at their ponds of origin.

We combined two temperature treatments (19 °C or 30 °C) with two hormone treatments (0 or 30 ng/L EE2) and applied their combinations over one of three treatment periods (Fig. 1). We replicated each treatment combination (temperature × EE2 × period) four times per family, resulting in 576 experimental animals. We chose the nominal concentration of 30 ng/L EE2 for two reasons. First, this value is environmentally realistic; for example, the average EE2 concentration in large rivers in Hungary is 0.14 ng/L (Avar et al., 2016), but higher concentrations up to 98.33 ng/L occur at point sources of pollution (Jakab et al., 2020). Second, a similar concentration (0.09 nM, equivalent to 27 ng/L) was shown to cause female-biased sex ratios in the common frog (*Rana temporaria*), a close relative of agile frogs (Pettersson and Berg, 2007). We chose a 30 °C temperature treatment because it represents an environmentally relevant extreme that occurs in the ponds where we collected eggs for this experiment (Szederkényi et al., unpublished data) and in other water bodies where amphibians breed and develop in climates similar to ours (Lambert et al., 2018; Lindauer et al., 2020).

**Fig 1.**
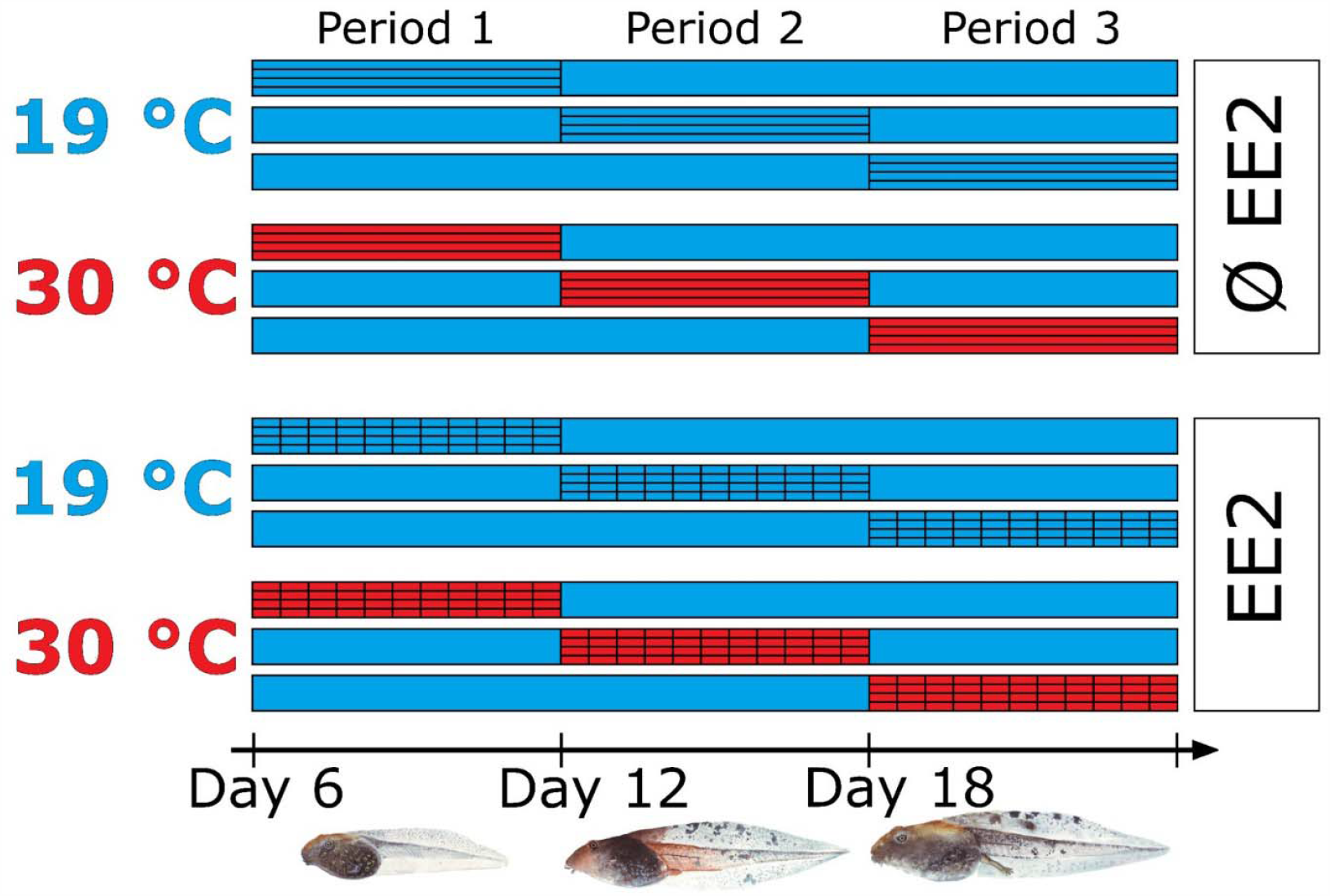
Schematic illustration of the 12 treatments, each horizontal bar representing a treatment group. Treatment periods are symbolized with horizontally striped bars; vertical stripes symbolize hormone treatment, and red bars symbolize high-temperature treatment.

Outside the treatment period, we reared tadpoles at 20.1 ± 1.1 °C (mean ± SD) individually in 2-L plastic containers filled with 1 L RSW, arranged in a randomized block design where each block contained all members of one family. We changed the rearing water two times a week, and fed tadpoles *ad libitum* with slightly boiled chopped spinach. The light:dark cycle was adjusted weekly to outdoor conditions, starting with 12:12 h in late March which we gradually changed to 14:10 h by the end of April. Each tadpole was exposed to a treatment period of six days, starting either 2, 8, or 14 days after start of the experiment (period 1: developmental stages 25-28, period 2: developmental stages 29-32, period 3: developmental stages 33-37). During the treatment period, treated tadpoles experienced the following changes in rearing conditions. The volume of rearing water was increased to 1.7 L, and the container was placed in an 80 × 60 × 12 cm tray filled with ca. 18 L tap water (for details see Supplementary material). The tap water was circulated using a Tetra WP 300 aquarium water pump and heated using a Tetra HT 300 aquarium heater. This arrangement ensured homogeneous water temperatures in the tray and among the tadpole rearing containers. Each tray hosted twelve containers, one from each family, resulting in four trays in each treatment group. During the treatment period we changed the water of treated animals every other day with temperature-adjusted and chemical-administered RSW (according to experimental treatments) and fed tadpoles with a reduced amount of spinach (for details see Supplementary material) to prevent water fouling in the heat treatments. During treatment, the water of all treated tadpoles contained a minute amount of ethanol (1 µL 96 % ethanol in 100 L RSW) because this solvent was necessary for EE2 treatment. This ethanol concentration is much lower than those that were shown to damage tadpoles (Fainsod and Kot-Leibovich, 2018; Peng et al., 2005; Taylor and Brundage, 2013) and in a previous experiment we showed that the same ethanol concentration did not result in skewed sex ratio or in any gonadal abnormalities in agile frogs (Bókony et al., 2020).

At the beginning of each treatment period we filled the trays with 19 °C water and placed the tadpoles’ containers (with freshly changed RSW) in the trays. In case of treatments involving elevated temperature, we subsequently turned on the heating. Thereby, water temperature gradually increased to 30 °C over the course of two hours and was maintained at 29.9 ± 0.2 °C (mean ± SD) in tadpoles’ containers. After six days of treatment, we turned off the heating in the trays, replaced the water in the rearing containers with 1 L fresh, 30 °C RSW, and transferred them back to their original place in the laboratory. These procedures ensured that the tadpoles were not exposed to a sudden change in temperature. In the 19 °C treatments, we applied the same protocol, except that the water in the trays was not heated, only circulated; water temperature in these treatments was 18.8 ± 0.3 °C (mean ± SD) in the tadpoles’ containers (for more details see Supplementary material).

To expose tadpoles to EE2 we used an analytical standard obtained from Sigma (E4876). The stock solution was prepared by dissolving 30 mg EE2 in 10 mL 96 % ethanol. At the beginning and at each water change during the treatment period, we added 1 µL of the stock solution into 100 L RSW, resulting in 30 ng/L EE2 in the water of hormone-treated tadpoles (and yielding the same ethanol concentration as in the water of tadpoles not treated with EE2). To determine how EE2 concentration changed during treatment, we took samples from the rearing water of treated tadpoles three times during the experiment. Samples were collected from each EE2 treatment (three samples from 19 °C, and another three from 30 °C) right before water change (i.e. two days after the nominal concentration was applied). One liter of each sample was collected into an amber PET flask, and transported to the analytical laboratory of the Centre for Agricultural Research in Martonvásár, where they were frozen and stored at −80 °C. Subsequent UPLC-UniSpray(tm) MS/MS analysis (for detailed methods see Bókony et al., 2021) showed that, over two days of treatment, the initial concentration has approximately halved at 19 °C (measured concentrations: 14.27, 15.93, 16.6 ng/L), and dropped to about a quarter at 30 °C (6.42, 7.53, 7.99 ng/L). We did not take samples from the water of tadpoles that were not treated with EE2, because our previous experiment showed that there is no EE2 contamination in the RSW we use in our laboratory (Ujhegyi and Bókony, 2020).

By starting the treatments 6, 12, or 18 days after hatching, we aimed to cover the majority of the larval period without risking that tadpoles would start to metamorphose during treatment, because a preliminary experiment showed that exposure to 30 °C during late tadpole development may result in increased mortality in agile frogs (unpublished data). We had no prior information on whether and when agile frogs have a sensitive time window for sex determination, but studies on other closely related species indicated such a time window either during early (Hogan et al., 2008) or mid-late larval development (Lambert et al., 2018). When tadpoles approached metamorphosis, we checked the rearing containers daily. When an individual reached developmental stage 42 (emergence of forelimbs), we noted the time until metamorphosis and measured body mass using a laboratory scale (± 0.1 mg). We changed the water volume to 0.1 L in the respective rearing container, slightly tilted it to prevent drowning, and covered it with a perforated, transparent lid to prevent escape. When metamorphosis was complete (developmental stage 46, disappearance of the tail), we moved the animal into a new rearing container, lined with wet paper towels as a substrate and a piece of egg carton as a shelter, which we changed every other week. Froglets were fed *ad libitum* with springtails and small (2-3 mm) crickets sprinkled with a 3:1 mixture of Reptiland 76280 (Trixie Heimtierbedarf GmbH & Co. KG, Tarp, Germany) and Promotor 43 (Laboratorios Calier S.A., Barcelona, Spain) containing vitamins, minerals and amino-acids.

We dissected the animals 6-8 weeks after metamorphosis (14-16 weeks after they reached the free-swimming tadpole stage). At this age the gonads are well differentiated in this species (Bernabò et al., 2011; Ogielska and Kotusz, 2004). When a froglet reached this age, we measured its body mass to the nearest 0.01 g and euthanized it using a water bath containing 6.6 g/L MS-222 buffered to neutral pH with the same amount of Na_2_HPO_4_. After dissection, we cut out the entire digestive tract and measured its mass to the nearest 0.01 g, because many animals’ guts contained food although we had not fed them for 2-4 days before dissection. We examined the gonads and the associated fat bodies under an Olympus SZX12 stereomicroscope at 16× magnification (Fig. S1). We recorded whether the animal had fat bodies and categorized phenotypic sex as male (testes), female (ovaries), or uncertain (abnormally looking gonads). We carefully removed the gonads and fixed them in neutral-buffered 10 % formalin (Sigma 1.00496) for histological analysis. We took a tissue sample (hind feet) from each froglet and stored it in 96 % ethanol until DNA extraction.

### 2.2. Histology

In a preliminary study we observed that sex categorized based on gonadal anatomy matched sex categorized by histology in all of 32 agile frogs (17 males, 15 females) that had been raised without any chemical treatment at 19 °C (Nemesházi et al., 2020a). Furthermore, in a previous study we found that spontaneous female-to-male sex reversal in agile frogs resulted in normal testicular histology, except for two masculinized individuals that had normal testis anatomy and histology with a single oogonium detected in each (Nemesházi et al., 2020a). Therefore, to minimize the costs of histological analysis in the present study, we chose to analyze gonad histology only in those froglets for which phenotypic sex was ambiguous based on gonad anatomy (Fig. S1). In these cases, when phenotypic sex could be clearly identified by histology, the animal was considered as female or male; if mixed-sex tissues were found, the individual was considered as intersex.

Histological sections were prepared at the Department of Pathology, University of Veterinary Medicine Budapest. The fixed gonads were placed in embedding cassettes and dehydrated through graded ethanol, cleared in xylene and infiltrated with paraffin wax in an Excelsior ES Tissue Processor (Thermo Fisher Scientific). The processed gonads were embedded in paraffin, sectioned into 3-4 μm longitudinal slices using a Reichert type microtome, stained with haematoxylin and eosin, and mounted on glass slides. The slides were examined and photographed using a Zeiss Axioskop 2 microscope equipped with an AxioCam ICc5 camera. For each individual, we examined 5-6 sections; ovaries were identified by the presence of diplotene oocytes, and testes by seminiferous tubules and spermatogonia (Fig. S2).

### 2.3. Genetic sexing

DNA was extracted using Geneaid Genomic DNA Extraction Kit for animal tissue (Thermo Fisher Scientific) following the manufacturer’s protocol, except that digestion time was 2 hours. Genetic sexing was performed using a recently developed molecular marker set which has been validated for agile frog populations in our study area (Nemesházi et al., 2020a). In short, we first tested all froglets for marker Rds3 (≥ 95 % sex linkage) applying high-resolution melting (HRM) and accepted an individual to be concordant male or female if its Rds3 genotype was in accordance with its phenotypic sex. Individuals that appeared to be sex-reversed based on Rds3 were tested for marker Rds1 (≥ 89 % sex linkage) using PCR, and were accepted to be sex-reversed only if both markers confirmed sex reversal. Individuals with discrepant genotyping results (i.e. contradiction between Rds1 and Rds3) were considered to be of unknown genetic sex. All details of HRM and PCR methods are described in Nemesházi et al., (2020a).

### 2.4. Statistical analyses

We analyzed the effects of treatments using pre-planned comparisons (Ruxton and Beauchamp, 2008). For each dependent variable, we ran a model (see model specifications below) containing the three-way interaction of temperature treatment, EE2 treatment, and treatment period; block (family) was entered as random factor in each model. Then, using the model estimates we tested the overall effect of temperature, EE2, and their interaction within each treatment period by calculating *a priori* linear contrasts (Lenth et al., 2020; Ruxton and Beauchamp, 2008). We evaluated the significance of linear contrasts both with and without correction for multiple testing; correction was applied by using the false discovery rate (FDR) method (Pike, 2011). All analyses were conducted in ‘R’ (version 3.6.3; R Core Team, 2020), with the ‘emmeans’ function of ‘emmeans’ package for linear contrasts. Sample sizes are given in Table 1.

**Table 1:** Number of animals that died or survived, percentage of animals that had fat bodies, genetic and phenotypic sex ratio (% males in sexable animals), and female-to-male sex-reversal rate (% of phenotypic males in genetic females) in each treatment group. Asterisks mark sex ratios that differ significantly from 1:1 according to binomial tests.

| Treatment group | Died before treatment | Died during treatment | Died after treatment | Died during metamorphosis | Died after metamorphosis | Dissected | Fat body % | Genetic sex ratio (% male) | Phenotypic sex ratio (% male) | Female-to-male sex reversal (%) |
| --- | --- | --- | --- | --- | --- | --- | --- | --- | --- | --- |
| Period 1 |  |  |  |  |  |  |  |  |  |  |
| Control | 0 | 1 | 3 | 2 | 2 | 40 | 86.8 | 55.0 | 55.0 | 0 |
| Heat | 1 | 26 | 1 | 0 | 0 | 19 | 52.6 | 63.2 | 89.5* | 71.4 |
| EE2 | 0 | 1 | 0 | 3 | 0 | 44 | 95.5 | 50.0 | 52.3 | 4.5 |
| Heat + EE2 | 3 | 21 | 2 | 1 | 0 | 21 | 61.9 | 78.9* | 94.7* | 75.0 |
| Period 2 |  |  |  |  |  |  |  |  |  |  |
| Control | 0 | 1 | 1 | 1 | 0 | 45 | 93.2 | 48.9 | 48.9 | 0 |
| Heat | 4 | 4 | 1 | 6 | 1 | 32 | 71.9 | 56.7 | 83.3* | 61.5 |
| EE2 | 2 | 0 | 0 | 3 | 0 | 43 | 95.4 | 53.5 | 55.8 | 5.0 |
| Heat + EE2 | 0 | 5 | 5 | 3 | 1 | 34 | 64.7 | 61.8 | 73.5* | 30.8 |
| Period 3 |  |  |  |  |  |  |  |  |  |  |
| Control | 1 | 0 | 1 | 5 | 0 | 41 | 92.7 | 46.3 | 48.8 | 4.5 |
| Heat | 0 | 4 | 1 | 5 | 1 | 37 | 78.4 | 58.3 | 100* | 100 |
| EE2 | 4 | 0 | 0 | 2 | 1 | 41 | 92.7 | 47.5 | 47.5 | 0 |
| Heat + EE2 | 1 | 5 | 1 | 3 | 0 | 38 | 73.7 | 38.2 | 97.1* | 95.2 |

For the analysis of survival, we used Cox’s proportional hazards model (‘coxme’ function of the ‘coxme’ package). Survival was categorized as 1: died during treatment, 2: died after treatment, but before the start of metamorphosis, 3: died during metamorphosis, 4: died after metamorphosis, before dissection, 5: the froglet survived until dissection. Animals that died before their respective treatment period (16 individuals, see Table 1) were left out of the analysis of survival. We entered survival as the dependent variable, and treated category 5 as censored observations (i.e. individuals that did not die before dissection were neither excluded nor treated as dead; instead, they were included in the analysis with the information that they were last observed alive after the individuals in category 4 died).

To analyze time to metamorphosis, body mass at metamorphosis, and body mass at dissection, we used linear mixed-effects models (‘lme’ function of the ‘nlme’ package). In the analysis of time to metamorphosis, we also allowed the variances to differ among treatment groups using the ‘varIdent’ function, because graphical model diagnostics indicated heterogeneous variances. When analyzing body mass at dissection, we subtracted gut mass from total body mass to provide a measure for lean body mass (i.e. without food remains), and we included the individual’s age (number of days from finishing metamorphosis until dissection) as a covariate and the combination of genetic and phenotypic sex (i.e. a 3-category factor: concordant male, concordant female, or sex-reversed individual) as a fixed factor. From the latter analysis, we excluded seven individuals for which genetic sex was unknown and three individuals that had intersex gonads.

For the analyses of the presence/absence of fat bodies and phenotypic sex ratio, we used generalized linear mixed-effects models with binomial error distribution and a logit-link function (‘glmmPQL’ function of the ‘MASS’ package). The seven individuals with unknown genetic sex and the three with intersex gonads were excluded from both models. In the model of fat bodies, the combination of genetic and phenotypic sex (concordant male, concordant female, or sex-reversed individual) was added as fixed factor, and age of the froglets was included as a covariate.

For the analysis of sex reversal, we could not apply the same modeling framework as for sex ratio, because of separation (i.e. the number of sex-reversed individuals was zero in certain treatment groups). Therefore, we used Firth’s bias-reduced logistic regression (‘logistf’ function of the ‘logistf’ package), which yields less biased estimates when separation is present in the data as compared to logistic regression. Because this method does not accommodate random effects and linear contrasts, we could not include block as a random factor, and we obtained our pre-planned comparisons as the model estimates from three separate analyses, one for each of the three treatment periods. We restricted this analysis to genetically female individuals, and the dependent variable was phenotypic sex, i.e. whether or not the individual was sex-reversed.

## 3. Results

### 3.1. Sex

Regardless of treatment timing, phenotypic sex ratio and sex-reversal rate were both significantly affected by heat treatment, whereas EE2 and its interaction with heat had no significant effects (Tables 1-4). Heat treatment significantly increased the proportion of phenotypic males by inducing sex reversal in genetically female individuals (Tables 1-4, Fig. 2): overall, 58 out of 199 genetically female individuals became phenotypic males, 55 of which received heat treatment. This masculinizing effect was strongest if heat was applied in the third period and weakest if it was applied in the second period (Tables 1, 3, 4). Additionally, the genetic sex ratio of froglets surviving to dissection was also male-biased for animals that received heat treatment in the first period (11 females to 27 males, binomial test: *P* = 0.014); all other treatment groups had genetic sex ratios close to 1:1 (Table 1). Furthermore, we found three intersex individuals which came from different treatment groups: one received heat but no EE2 in the second period, one received EE2 but no heat in the third period, and one received both EE2 and heat in the third period. These animals were genetic females with abnormally looking gonads (Fig. S1) that were identified as ovotestes in the histological analysis (Fig. S2). We could not identify the genetic sex of seven phenotypically male individuals that received heat treatment, because in all these cases, the Rds3 genotype (corresponding to the locus with the highest sex linkage in our marker set) indicated sex reversal from female to male, whereas the Rds1 genotype suggested the genetic sex to be male.

**Table 2.**
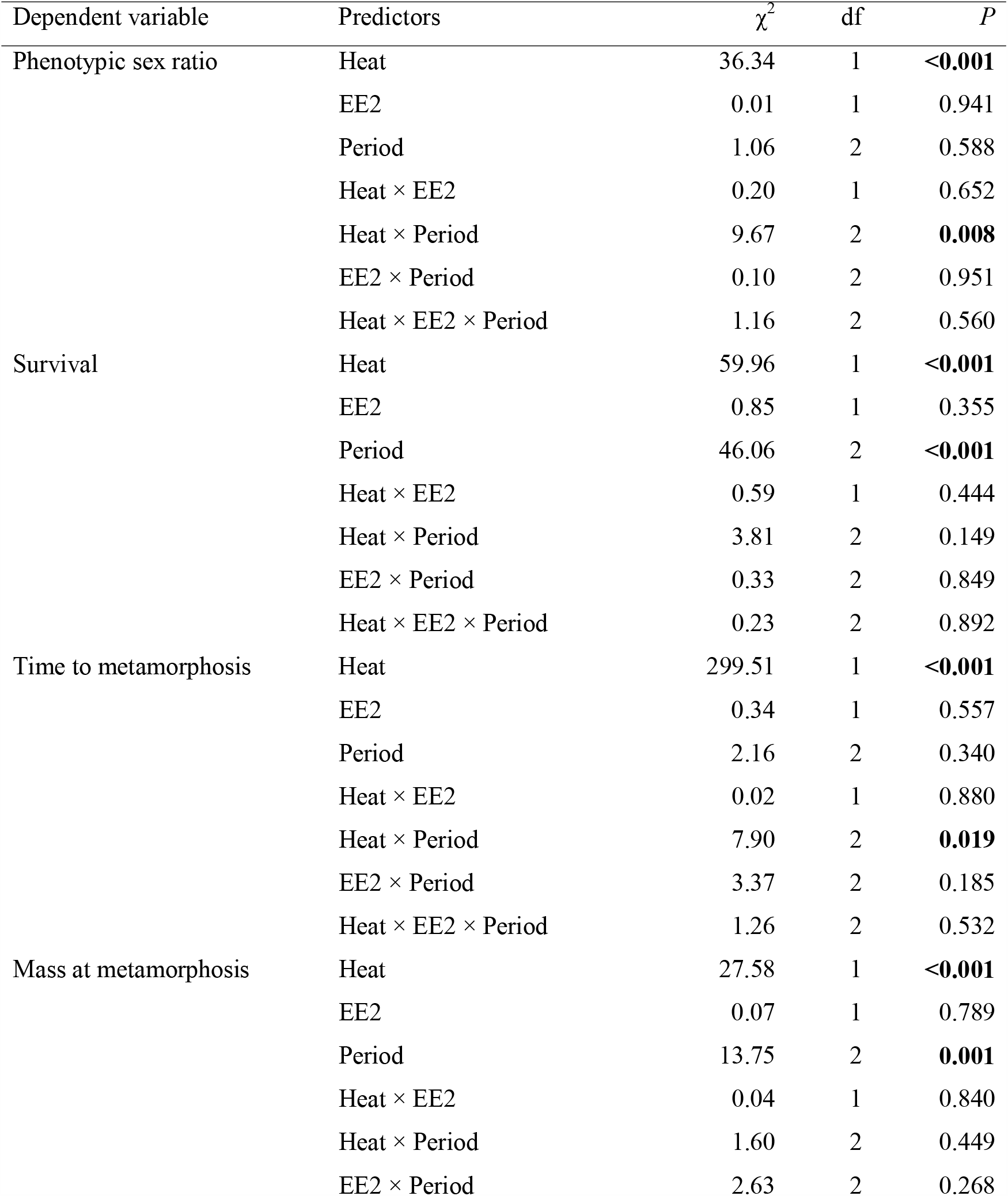
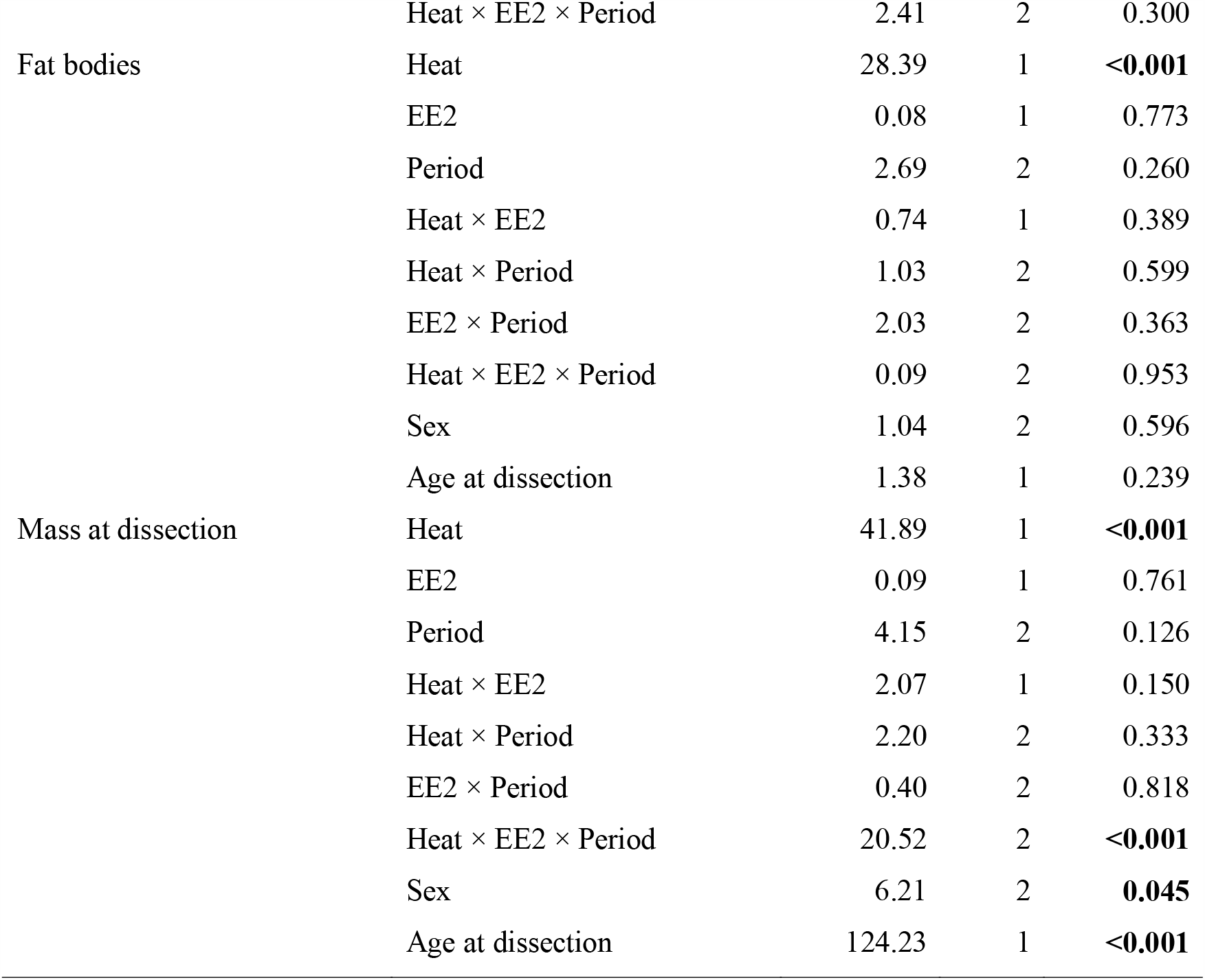
Type-2 analysis-of-deviance tables of the statistical models. Significant effects (*P* < 0.05) are highlighted in bold.

**Table 3.**
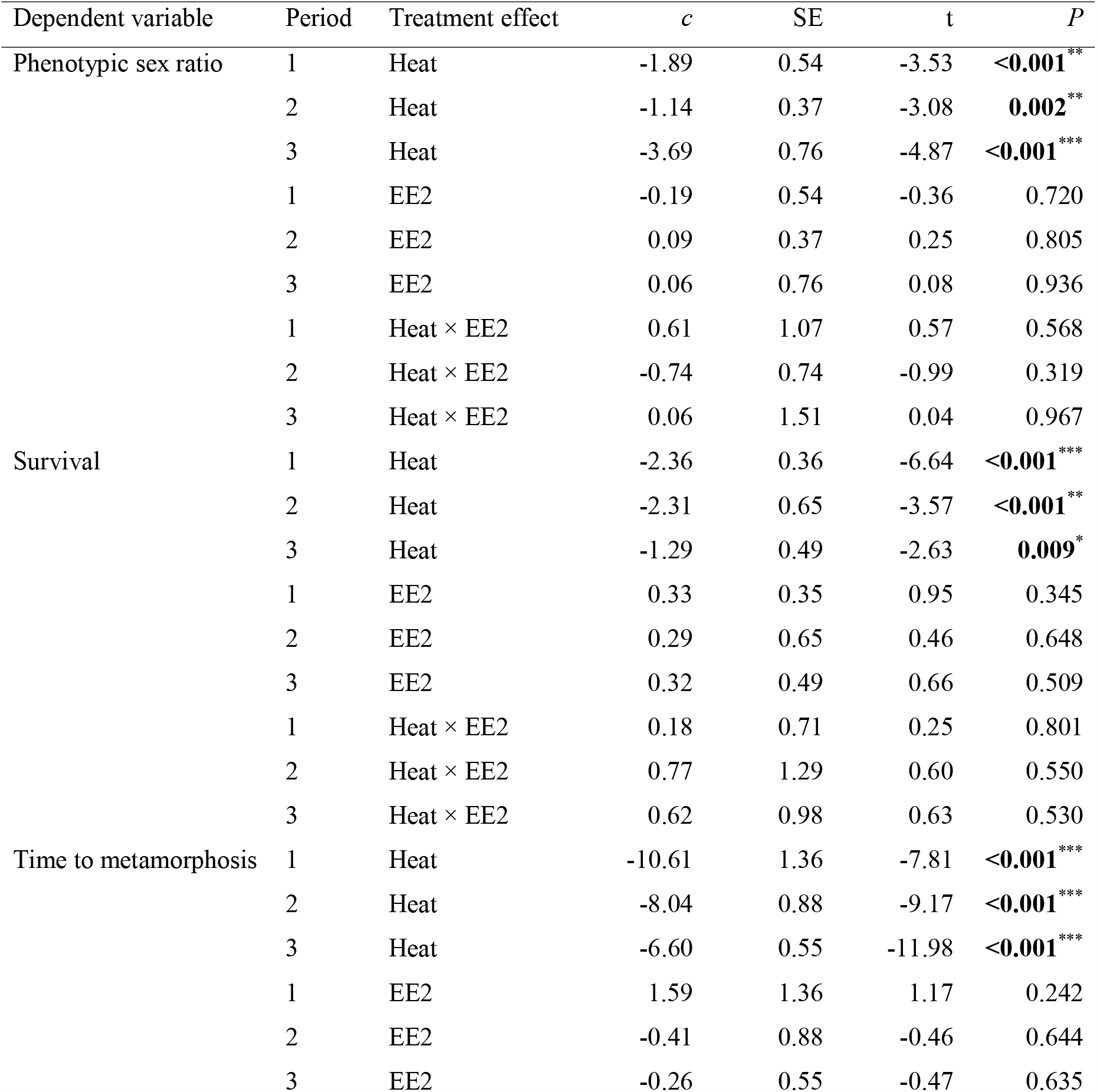
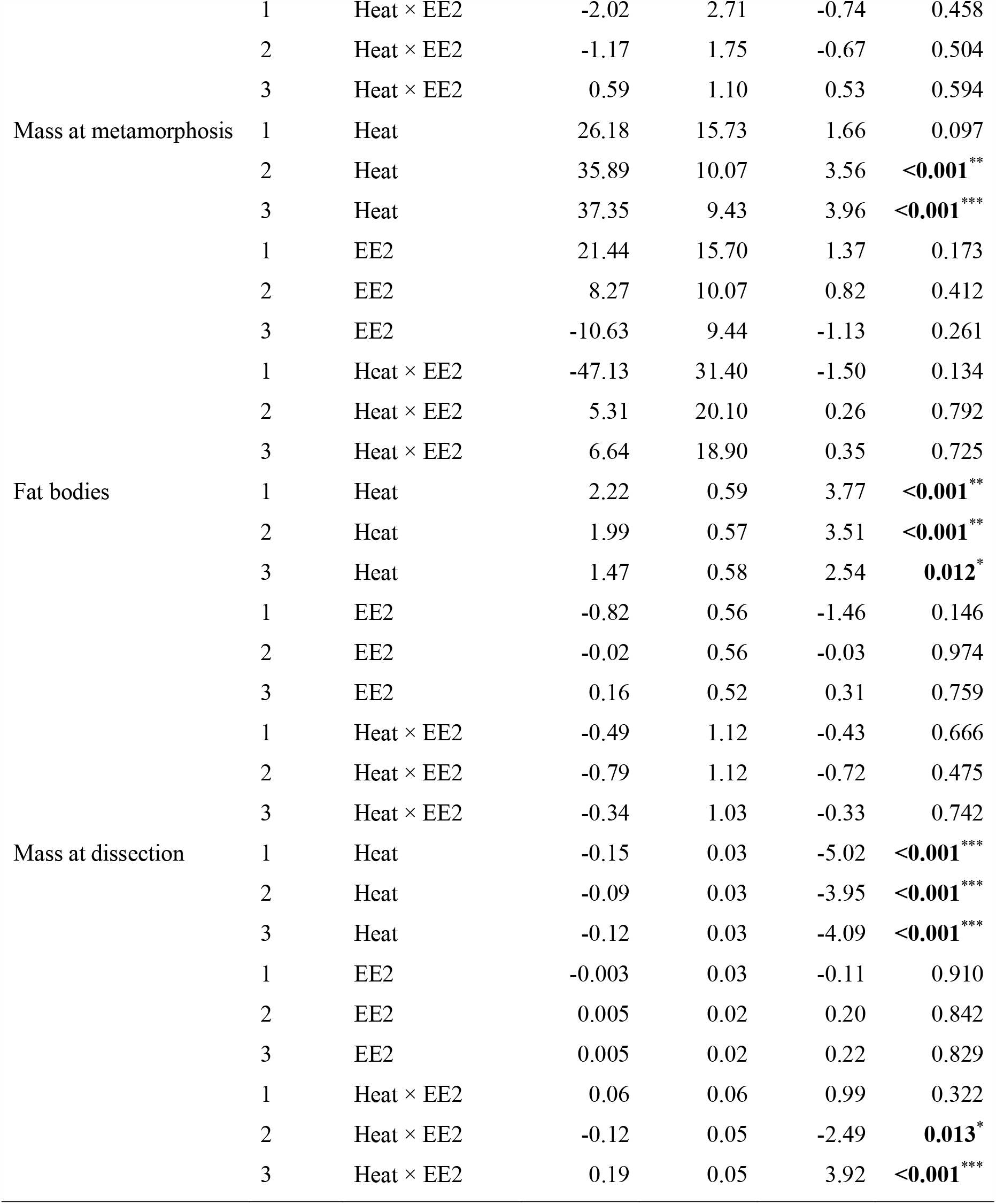
Results of pre-planned comparisons from the models in Table 2. Linear contrasts (*c*) with standard errors (SE) are reported with non-adjusted *P*-values; asterisks indicate if the results remain significant after FDR correction performed separately for each dependent variable (^*^*P* = 0.01 – 0.05, ^**^*P* = 0.001 – 0.01, ^***^*P* < 0.001).

**Table 4.**
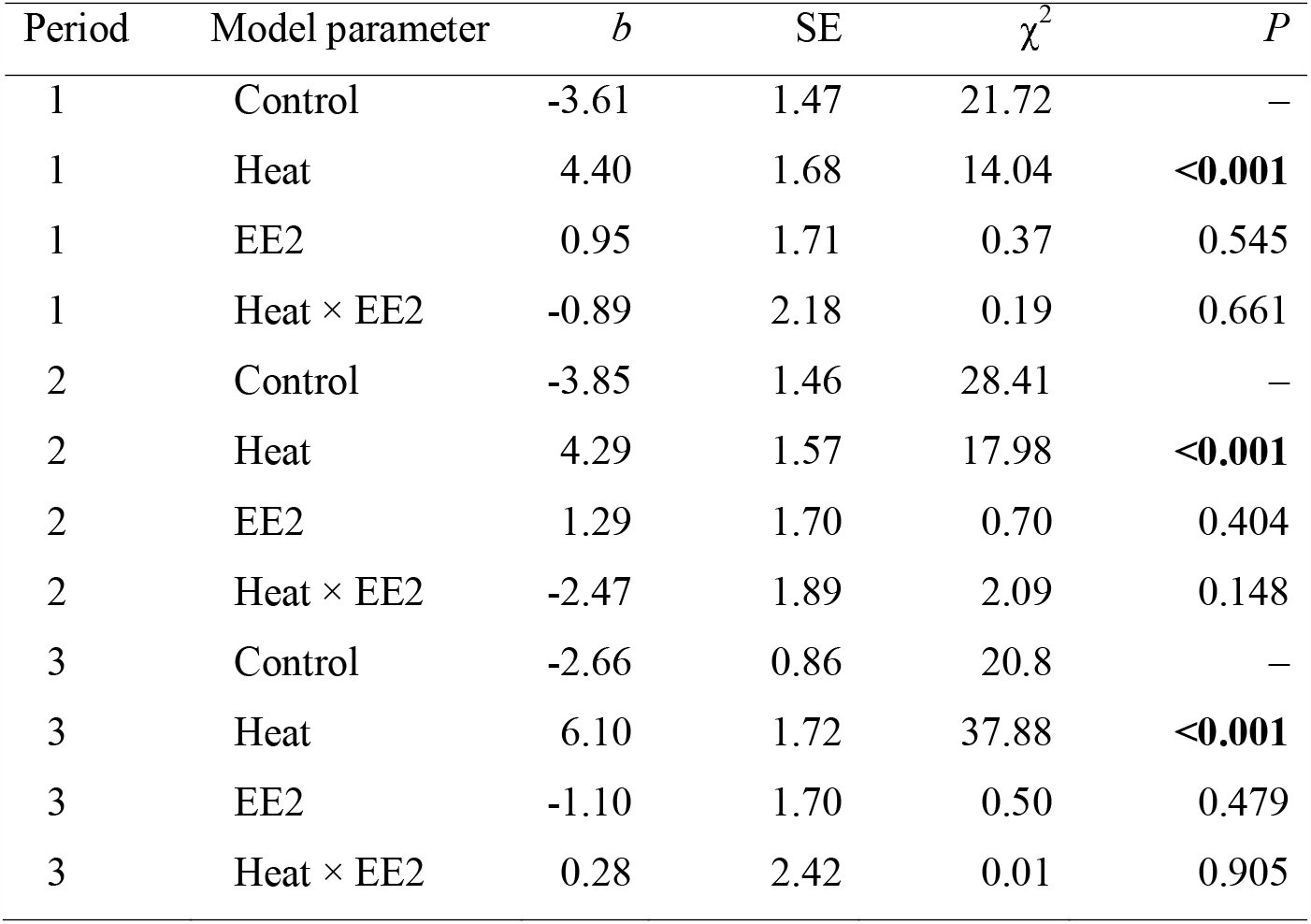
Results of Firth’s bias-reduced logistic regression models on female-to-male sex reversal. Parameter estimates (*b*) with standard errors (SE) are given on logit scale. In each period, the first parameter refers to the mean of the control group (19 °C, no EE2), whereas the second and third parameters refer to the effect of heat without EE2 and the effect of EE2 without heat, respectively. The fourth parameter refers to the interaction, i.e. the effect of EE2 on the effect of heat. We report *P* values only for parameters that refer to differences between treatments.

**Fig 2.**
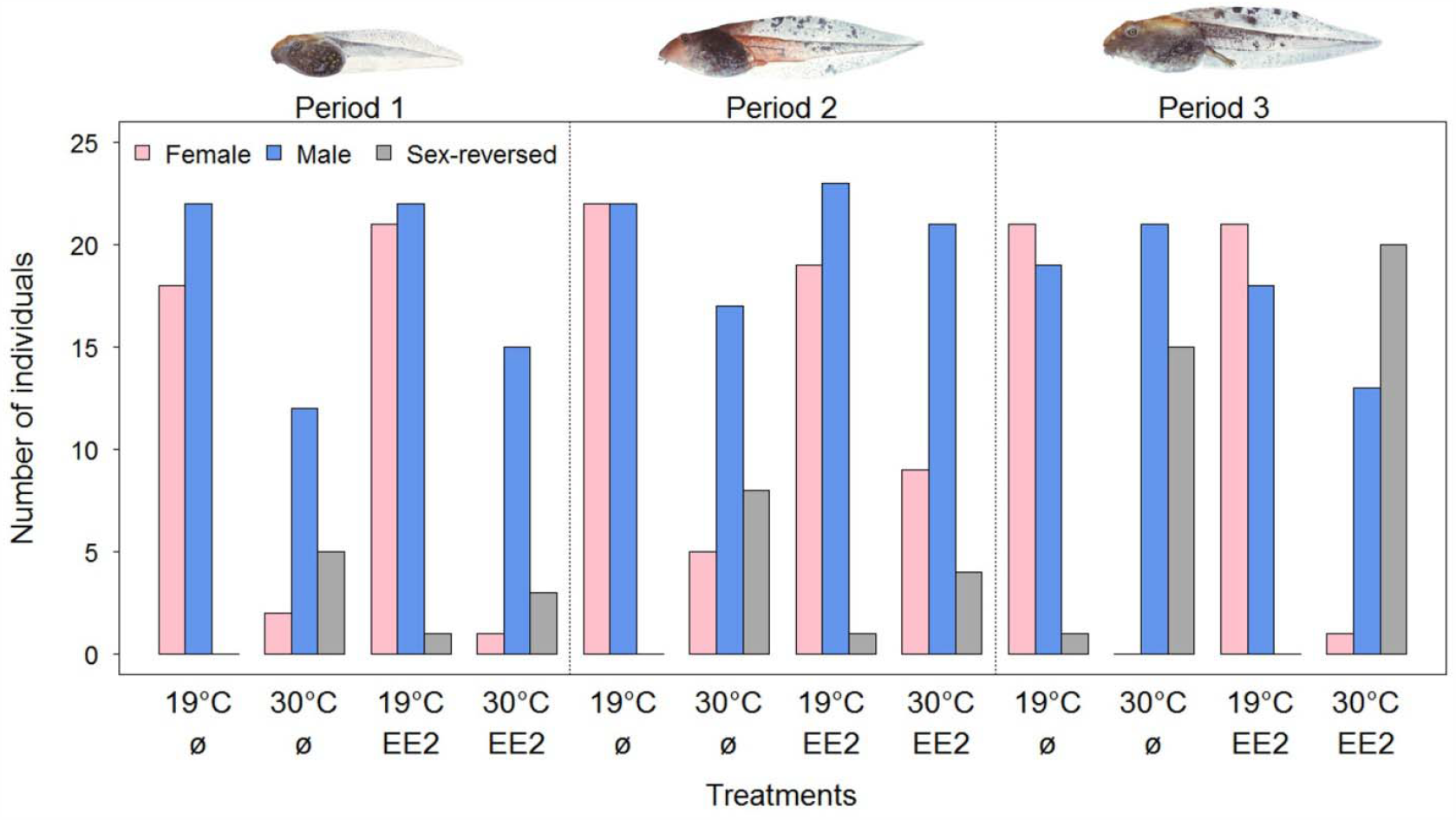
Effects of heat (30°C *versus* 19°C) and EE2 (30 ng/L, marked as EE2, *versus* none, marked with a Ø symbol) treatments on phenotypic sex ratio and female-to-male sex-reversal.

### 3.2. Survival, metamorphosis, and growth

In each of the three treatment periods, all dependent variables were significantly affected by heat (Tables 1-3), whereas EE2 and its interaction with heat had no significant effects (Tables 1-3) except for juvenile body mass (Table 2). Heat treatment significantly decreased survival (Tables 2-3), delayed metamorphosis (Tables 2-3, Fig. 3), decreased body mass at metamorphosis (Tables 2-3, Fig. 3), and increased the proportion of animals that had no fat bodies (Tables 1-3). Most of these effects were similar across the three treatment periods, except for the time to metamorphosis, which was delayed significantly more if heat was applied to younger tadpoles (Tables 2-3). For body mass of froglets at dissection, besides the main effect of heat, the three-way interaction (heat × EE2 × period) was also significant (Table 2). Heat applied in the first period increased froglet mass regardless of EE2 treatment, but for the other two periods the heat × EE2 interaction was significant (Table 3, Fig. 3). Without heat, EE2 applied in the second period tended to increase body mass, whereas EE2 applied in the third period decreased mass, but these effects were reversed in the heat-treated groups (Fig. 3). Thus, only the chemically untreated (no EE2) animals in the second period and the EE2-treated animals in the third period showed significant increase in body mass in response to heat (Fig. 3). Additionally, the combination of genetic and phenotypic sex also affected froglet mass significantly (Table 2): sex reversal reduced body mass by 5 % on average, to 1.15 ± 0.03 g compared to 1.22 ± 0.02 g in concordant females and 1.19 ± 0.02 g in concordant males (Table 5) across all treatment groups (Fig. 4).

**Table 5.** Differences in body mass (g) at dissection between female-to-male sex-reversed individuals, concordant males, and concordant females. Linear contrasts (*c*) with standard errors (SE) are reported with non-adjusted *P*-values, calculated from the model in Table 2. Asterisk indicates if the result remained significant after FDR correction (^*^*P* = 0.01 – 0.05).

| Linear contrasts | *c* | SE | t | *P* |
| --- | --- | --- | --- | --- |
| Concordant female vs. concordant male | 0.03 | 0.02 | 1.69 | 0.09 |
| Sex-reversed female vs. concordant female | -0.07 | 0.03 | -2.43 | **0.02\*** |
| Sex-reversed female vs. concordant male | -0.04 | 0.03 | -1.68 | 0.09 |

**Fig 3.**
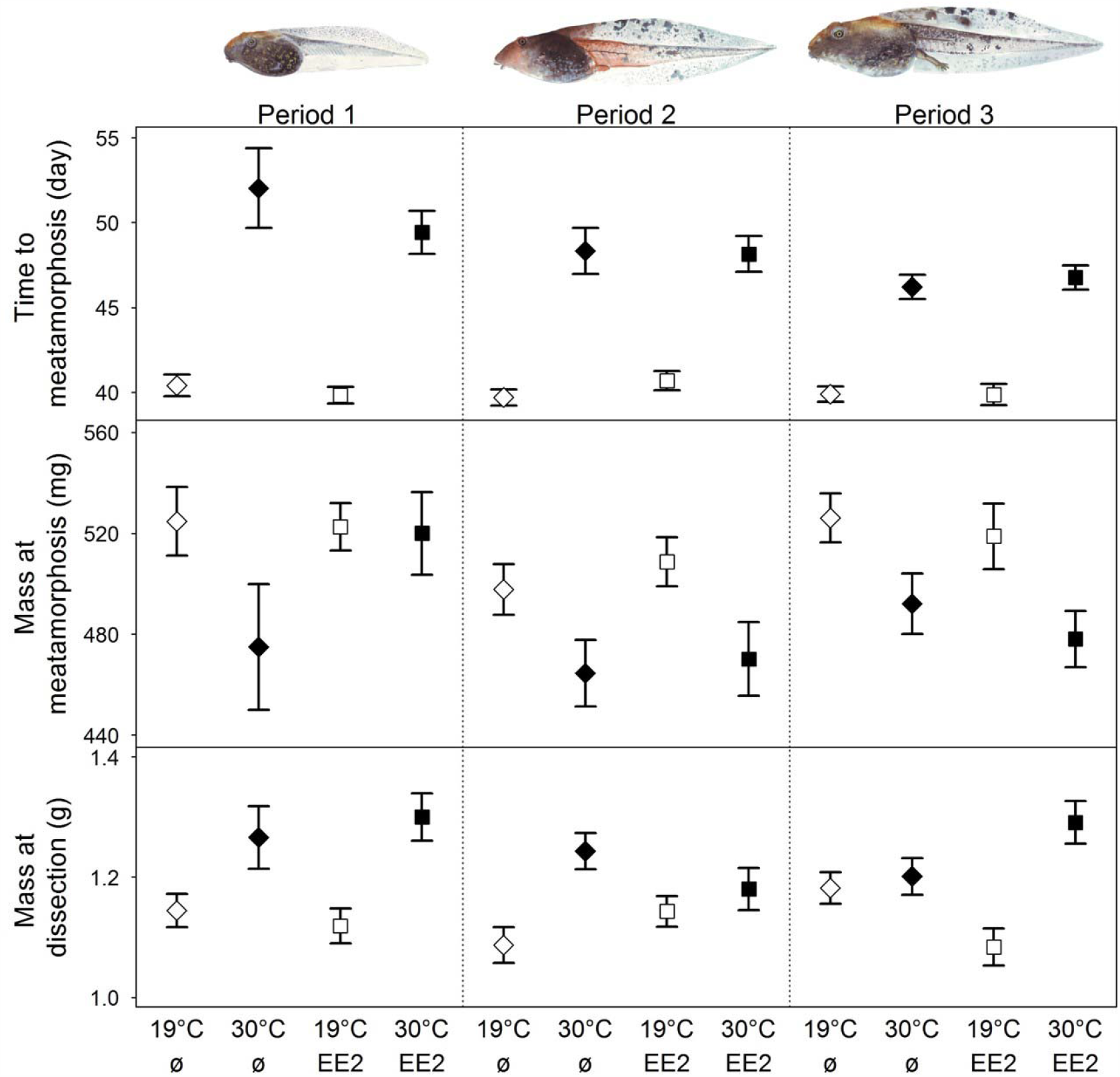
Effects of heat (30°C *versus* 19°C) and EE2 (30 ng/L, marked as EE2, *versus* none, marked with a Ø symbol) treatments on larval developmental time, body mass at metamorphosis, and the body mass of juvenile frogs at dissection. Error bars show the means and standard errors estimated from the models in Table 2.

**Fig 4.**
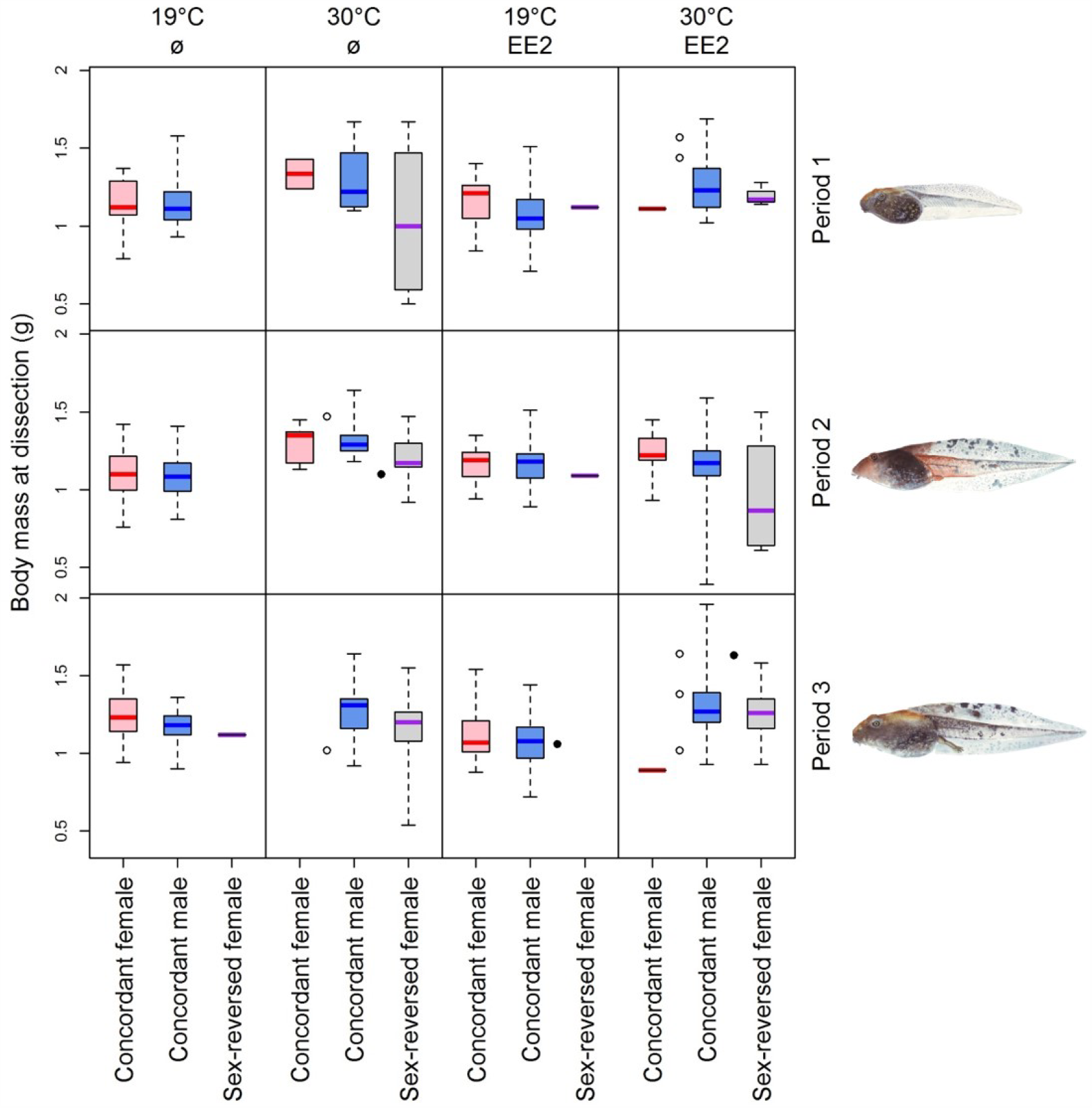
Boxplots (medians, interquartile ranges, and data ranges) of juvenile body mass for concordant females, concordant males, and female-to-male sex-reversed individuals. Empty and filled circles mark phenotypic males with unknown genetic sex and intersex individuals, respectively. Treatment groups are indicated as in Figures 2-3.

## 4. Discussion

In this study, we investigated the simultaneous effects of ecologically relevant larval exposure to high temperature and EE2 on the sexual development, survival, growth and somatic development of the agile frog. We found that all measured variables were significantly affected by the heat treatment regardless of its timing, whereas EE2 and its interaction with heat had no significant effects except on froglet body mass. The simulated heat wave caused masculinization and had long-term negative effects on several fitness-related traits, suggesting that populations of the agile frog (and other species with similar sensitivity to heat) face multiple threats from climate change due to skewed sex ratios and poor survival, which are not countered by environmental concentrations of feminizing pollutants like EE2.

In our experiment, the lack of EE2 effects on sex was unexpected, given that a similar concentration (27 ng/L) caused female-biased sex ratios in the closely related common frog (Pettersson and Berg, 2007). This discrepancy confirms that EE2 sensitivity can vary not only among phylogenetically distant lineages (Tamschick et al., 2016) but also within a genus (Mackenzie et al., 2003). A notable difference is that previous studies showing the feminizing effects of EE2 in amphibians either exposed individuals throughout their entire larval development and/or used doses much higher than the environmentally realistic concentrations (Berg et al., 2009; Gyllenhammar et al., 2009; Hogan et al., 2008; Mackenzie et al., 2003; Pettersson and Berg, 2007; Tompsett et al., 2013, 2012). Our treatments were ecologically relevant in terms of both magnitude and duration, as EE2 typically occurs in surface waters in the ng/L concentration range and its presence is usually not constant, possibly due to photolysis and adsorption to suspended solids and sediment (Avar et al., 2016; Bhandari et al., 2015; Jakab et al., 2020; National Center for Biotechnology Information, 2020). Therefore, our findings suggest that population persistence in the agile frog (and other species with similar sensitivities) is threatened more by climate change than by xenoestrogens.

Our experimental simulation of a six-days heat wave confirmed that high temperatures during early ontogeny can result in male-biased sex ratios, which was found for several other amphibian and fish species in previous studies. However, these earlier experiments usually applied heat treatment throughout the entire larval period (Chardard et al., 2004; Dournon et al., 1990; Eggert, 2004; Flament, 2016; Lambert et al., 2018; Ospina-Alvarez and Piferrer, 2008), which may not represent ecologically realistic temperature regimes in natural water bodies under current climatic conditions (Lambert et al., 2018; Lindauer et al., 2020). The bias towards phenotypic males in our experiment was mostly due to sex reversal, as 30-100 % of genetically female individuals (depending on treatment) developed testes after experiencing the heat weave. Thus, our study shows that even a relatively short hot spell, lasting only six days, can lead to a preponderance of males via sex reversal. Theoretical models suggest that such sex-ratio skews, due to increasing frequency of climate-driven masculinization, may have detrimental consequences for population viability (Bókony et al., 2017; Nemesházi et al., 2020b; Quinn et al., 2011; Schwanz et al., 2020; Wedekind, 2017). Additionally, it seems that the high temperature in our experiment caused sex-dependent mortality in the youngest tadpoles, because genetic sex ratio at dissection was also male-biased when heat was applied in the first period. This highlights the importance of molecular sexing methods for diagnosing sex reversals and disentangling the mechanisms by which anthropogenic disturbances cause skewed sex ratios, i.e. sex reversal vs. sex-biased mortality (Geffroy and Wedekind, 2020).

In many ectothermic vertebrates, there is a thermosensitive period (TSP), a time window of limited length during early development, during which environmental temperatures can influence whether the bipotent gonad commits to male or female development (Eggert, 2004; Flament, 2016; Mitchell and Janzen, 2010; Ospina-Alvarez and Piferrer, 2008). During the TSP the gonads are histologically undifferentiated, and the end of the TSP corresponds to the beginning of the meiotic prophase of ovarian differentiation (reviewed in Baroiller and Guiguen, 2001; Chardard et al., 2004). In the agile frog ovarian differentiation was reported to start at developmental (Gosner) stage 31 (Falconi et al., 2001), although gonad development in anurans may be tied to chronological age rather than to the stage of somatic development (Ogielska and Kotusz, 2004). In our experiment, tadpoles were between 6 and 24 days after hatching (developmental stages 25-37) during the treatment periods, and heat-induced sex-reversal occurred in all three 6-days periods. Thus, while our experiment does not allow for identifying the exact start and end of the TSP in agile frogs, it seems that it spans at least one quarter of their larval development. We propose that TSP length and timing deserve more attention in ectothermic vertebrates that are susceptible to sex reversal, because longer and later-occurring TSP may increase the risk that it coincides with extreme temperatures (e.g. early-summer heat waves) and thereby causes sex reversal or gonadal abnormalities.

Notably, three individuals in our experiment, out of 253 that did not receive heat treatment, showed a mismatch between genetic and phenotypic sex. This may be explained by misdiagnosed genetic sex due to sex-chromosome recombination or mutation, which occur in amphibians and make genetic sexing difficult (Alho et al., 2010; Lambert et al., 2016; Stöck et al., 2013). Alternatively, sex reversal may have occurred naturally, which has been reported to take place occasionally in several species (Holleley et al., 2016; Lambert, 2015; Nemesházi et al., 2020a) due to random processes affecting sex determination (Perrin, 2016). With this latter interpretation, here we found a 1.2 % baseline rate of female-to-male sex reversal in agile frogs, which is similar to the 4.8 % found in a previous study on lab-raised agile frogs (Nemesházi et al., 2020a). In the wild, however, the frequency of female-to-male sex-reversed individuals was considerably higher, especially in anthropogenic habitats including urban ponds (Nemesházi et al., 2020a). These findings combined with our current experimental results suggest that anthropogenic stressors, such as heat waves exacerbated by the urban heat island effect, increase sex-reversal rates above their baseline in natural populations.

Our current study also provides indirect but much-needed information about the relationship between sex reversal and individual fitness prospects. The results of our experiment show that, when sex reversal is triggered by heat stress, it is accompanied by reduced survival, slower development, lower body mass at metamorphosis, and less fat before the first winter hibernation. These changes suggest that sex reversal is associated with inferior fitness in agile frogs, corroborating a previous study (Nemesházi et al., 2020a) in which a much smaller sample of spontaneously masculinized froglets showed several signs of poor condition and increased physiological stress. These findings are in line with the emerging view that sex reversal is mechanistically linked with physiological stress, as studies on fish and reptiles suggest that extreme temperature may be one out of many stressors that influence sex development *via* the activation of the hypothalamus-pituitary-interrenal axis (Castañeda Cortés et al., 2019; Fernandino et al., 2013) or cellular pathways of the calcium and redox regulation system (Castelli et al., 2020). However, empirical data on the relative fitness of sex-reversed individuals in nature are so far few and controversial (Holleley et al., 2016; Senior et al., 2012); for example, despite the poor health of agile frogs masculinized in the laboratory, body mass of female-to-male sex-reversed adults did not differ from normal males’ in free-living populations (Nemesházi et al., 2020a).

In our current experiment, the negative effect of heat treatment on metamorphic size disappeared after ca. two months: body mass at dissection was larger in heat-treated animals than in their control siblings. This result might be explained with compensatory growth (Hector et al., 2012; Squires et al., 2010). Alternatively, the environmental matching hypothesis claims that individuals that developed under poor conditions can have higher fitness later in life than those that developed under good conditions, if adult environmental conditions are also poor (Monaghan, 2008). Whatever mechanism allowed for faster growth after metamorphosis, it pertained only to those heat-treated individuals that developed concordant sex; those that underwent masculinization had reduced body mass as juveniles. This suggests that heat-induced sex reversal constrains juvenile growth, or alternatively, individuals with the least inherent potential for juvenile growth performance are the most likely to respond to heat with sex reversal. This latter possibility might even be an adaptive strategy if reduced growth is more detrimental for fitness in females than in males (Baroiller and D’Cotta, 2016; Schwanz et al., 2016), which seems likely in agile frogs and many other ectothermic vertebrates where mature females are larger than mature males.

Juvenile body mass was the only trait in our study that was affected by EE2, but only in some treatment combinations. When treatment was applied in the second period, EE2 counteracted the heat-induced increase in body mass, whereas in the third period, EE2 decreased mass without heat but contributed to mass increase when combined with heat. These results indicate that heat waves and EE2 pollution in water bodies may have non-additive effects on fitness via influencing body mass even when EE2 does not cause feminization in environmentally relevant concentrations. The mechanisms behind these interactions are unclear, but both high temperature and estrogens may influence the hormonal regulation of growth, including thyroid hormones, prolactin and growth hormone (Hogan et al., 2008; Hu et al., 2019). For example, a study on salamander larvae found that high temperature decreased the gene expression of growth hormone and its brain receptors during treatment but increased it after treatment (Hu et al., 2019). In amphibians, growth is governed by different hormones in different stages of development, which might explain the stage-dependent effects of EE2 (Hogan et al., 2008).

## 5. Conclusions

Taken together, our experiment showed that a six-days heat wave induces female-to-male sex reversal and reduces growth, development and survival, whereas exposure to environmentally relevant concentration of EE2 is unlikely to temper the masculinising effects of high temperature. Nevertheless, EE2 and heat had non-additive effects on juvenile body mass, and additionally, sex reversal was associated with reduced mass regardless of treatment. These results highlight that climate change and chemical pollution may have intricate consequences for individual fitness and population persistence in species with environmentally sensitive sex determination. Understanding these effects is essential for the conservation of biodiversity in our human-modified world.

## Supporting information

Electronic supplementary material

## Funding

The study was funded by the National Research, Development and Innovation Office of Hungary (NKFIH, grants 115402 to V.B. and 124375 to A.H.). The authors were supported by the János Bolyai Research Scholarship of the Hungarian Academy of Sciences (to V.B., A.H., and O.I.H.), the ÚNKP-20-5 New National Excellence Program of the Ministry for Innovation and Technology from the source of the National Research, Development and Innovation Fund (“Bolyai+ Scholarship” to V.B. and A.H.), the Ministry of Human Capacities (National Program for Talent of Hungary, NTP-NFTÖ-18-B-0412 to V.V., NTP-NFTÖ-17-B-0317 to E.N.), the Austrian Agency for International Cooperation in Education & Research (OeAD-GmbH; ICM-2019-13228 to E.N.), and the NKFIH (124708 to O.I.H.). None of the funding sources had any influence on the study design, collection, analysis, and interpretation of data, writing of the paper, or decision to submit it for publication.

All experimental procedures were approved by the Ethical Commission of the Plant Protection Institute and carried out according to the permits issued by the Government Agency of Pest County (Department of Environmental Protection and Nature Conservation, PE/KTF/3596□6/2016, PE/KTF/3596□7/2016 and PE/KTF/3596□8/2016), the Budapest Metropolitan Municipality (Department of City Administration, FPH061/2472-4/2017) and the EC Directive 86/609/EEC for animal experiments (http://europa.eu.int/scadplus/leg/en/s23000.htm).

## Acknowledgements

We are grateful to Kamirán Á. Hamow and Mihály Dernovics for the chemical analysis of water samples. We thank Árpád Ferincz, Ádám Staszny, Lilianna Olimpia Pap, Vera Juhász and András Weiperth for sharing their unpublished data on EE2 concentrations in Hungarian streams. We are grateful to all members of the Lendület Evolutionary Ecology Research Group for insightful discussions, and Márk Szederkényi, Eszter Nádai-Szabó, Csenge Kalina, Zsófia Boros, Boglárka Jaloveczki, Stephanie Orf and Gergely Tarján for help with animal care and data archiving. We thank Gergő Tholt and the NÖVI Department of Zoology for allowing us to use their stereomicroscope and camera and for providing helpful advice. We are grateful to Renáta Pop and the Department of Pathology, University of Veterinary Medicine Budapest for preparing histological sections, and to Beata Rozenblut-Koscisty and Maria Ogielska for help with interpreting histological images. We thank Bálint Bombay for the paintings of agile frog tadpoles.

## Competing interests

The authors declare that they have no known competing financial interests or personal relationships that could have appeared to influence the work reported in this paper.

## Author contributions

Conceptualization: V.B.; Methodology: Z.M., E.N., N.U., J.U., A.H., V.B.; Investigation: Z.M., E.N., N.U., V.V., J.U., A.K., R.B., N.V., Z.G., O.I.H, A.H., V.B.; Data curation: Z.M., R.B., V.B.; Formal analysis: Z.M., V.B.; Writing - original draft: Z.M., V.B.; Writing - review & editing: Z.M., E.N., N.U., V.V., J.U., A.K., R.B., N.V., Z.G., O.I.H, A.H., V.B.; Visualization: Z.M., V.B.; Supervision: V.B.; Funding acquisition: A.H., V.B.

## Highlights

- Experimentally simulated heat wave caused female-to-male sex reversal in tadpoles
- Exposure to ethinylestradiol (EE2) did not alter the masculinizing effect of heat
- Heat also reduced survival, development speed, and body mass at metamorphosis
- Heat and EE2 had non-additive effects on body mass two months post-metamorphosis
- Sex-reversed individuals had reduced body mass two months post-metamorphosis

## Notes

### Competing Interest Statement

The authors have declared no competing interest.

