## Supplementary material for "Sex reversal and ontogeny under climate change and chemical pollution: are there interactions between the effects of elevated temperature and a xenoestrogen on early development in agile frogs?": Electronic supplementary material

**Supplementary Methods**

*Measurements validating temperature in heat treatments*

Following the setup of trays (filled with ca. 18 L tap water, resulting in a water depth of 8 cm) to heat treatments, but before the start of animals’ exposure, we tested the heating system with boxes that were the same as individuals’ rearing containers (filled with 1.7 L RSW, resulting in a water depth of 10 cm) and measured the exact water temperature in containers of each position of the tray, plus the temperature of the circulated tap water. We replicated measurements ten times altogether on two consecutive days with a Greisinger digital thermometer (GTH175/PT). After the termination of the experiment, we repeated these measurements five times. In order to record incidental temperature fluctuations during treatment periods, we daily checked the temperature of tap water in the trays with the digital thermometer. Furthermore, automated data loggers (Onset HOBO Pendant Temperature/Light 8K Data Logger) also recorded water temperature in each tray every 30 minutes during the whole experiment. Accidental temperature fluctuations were not detected in the trays, and measurements in the same positions were very similar before and after the experiment. Therefore, we estimated the temperature likely experienced by individuals during the experiment from daily measured tray water temperature, correcting it with the average difference we measured between the water in each container (in each position in the trays) and the mean temperature of the water in the respective tray. This method minimized the disturbance caused to the animals by daily measurements, but delivered sufficient data to draw conclusions about the temperature experienced by the animals during treatment periods, and validated the operability of the applied heating setup.

*Feeding during the heating treatments*

During the treatment period tadpoles were fed as follows. We homogenized 80 g slightly boiled chopped spinach in 235 mL RSW with a hand blender, and added five drops in the first, seven drops in the second, or nine drops in the third treatment period to the containers (ca. 0.06 mL per drop) after every water change. We used 5 ml Pasteur pipettes, of which we cut off the last 2.8 cm from the tips.

**Fig. S1.** Gonads in juvenile agile frogs at 16× magnification: A) normal testes (t), B) normal ovaries (o), C) intersex gonads (ovotestes; ot); and varying amounts of fat bodies (f).

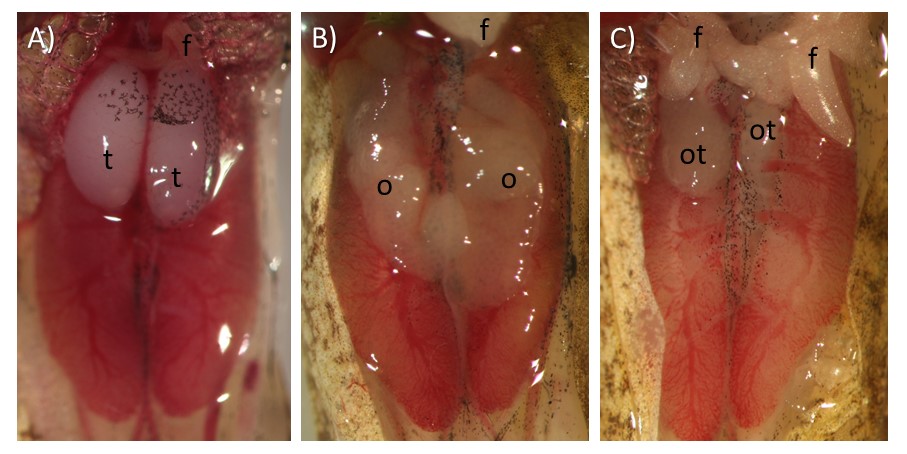

**Fig. S2.** Histological sections of agile frog gonads (10×): A) normal testis with seminiferous tubules (black arrowheads) and somatic cells forming the precursor of rete testis (yellow arrow), B) normal ovary with previtellogenic diplotene oocytes about 120-200 µm in diameter, C) ovotestis with seminiferous tubules, previtellogenic diplotene oocytes (o), and degenerating oocytes (red arrowheads). D) Larger magnification (20×) of an ovotestis, with somatic tissue forming a seminiferous tubule (encircled yellow) between normal and degenerating oocytes.

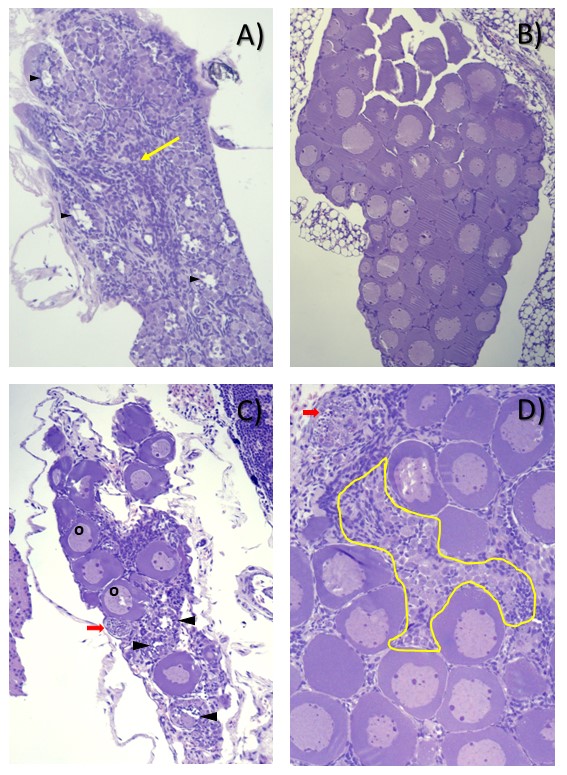
